# idCOV: a pipeline for quick clade identification of SARS-CoV-2 isolates

**DOI:** 10.1101/2020.10.08.330456

**Authors:** Xun Zhu, Ti-Cheng Chang, Richard Webby, Gang Wu

**Author notes:** The authors wish it to be known that these authors contributed equally (Xun Zhu, Ti-Cheng Chang).

## Abstract

idCOV is a phylogenetic pipeline for quickly identifying the clades of SARS-CoV-2 virus isolates from raw sequencing data based on a selected clade-defining marker list. Using a public dataset, we show that idCOV can make equivalent calls as annotated by Nextstrain.org on all three common clade systems using user uploaded FastQ files directly. Web and equivalent command-line interfaces are available. It can be deployed on any Linux environment, including personal computer, HPC and the cloud. The source code is available at https://github.com/xz-stjude/idcov. A documentation for installation can be found at https://github.com/xz-stjude/idcov/blob/master/README.md.

## Introduction

The on-going Coronavirus disease 2019 (Covid-19) pandemic has resulted in over 734,000 deaths, affecting more than 188 countries and territories (CSSE, 2020; Dong, et al., 2020). The general impact of the disease calls for a quick response from the local health system at every stage of its transmission.

Severe acute respiratory syndrome coronavirus 2 (SARS-CoV-2; previously known as 2019-nCoV) is the pathogenic cause of Covid-19 (Lescure, et al., 2020). Quickly identifying the genetic clade of the virus helps develop an understanding of the local transmission of the disease. The recent NYC study (Gonzalez-Reiche, et al., 2020), for example, identified crucial local transmission events, plurality of virus introductions, and the possible sources of these introductions. The classification of the infected virus can also provide more granular information to contact tracing and for epidemiological studies to understand disease severity and host genetic susceptibility to different lineages of SARS-CoV-2.

In order to quickly identify the clade of an isolate of SARS-Cov-2 given its sequencing FastQ files, we have developed a bioinformatics pipeline and a user-friendly web interface. An equivalent command-line interface is also supplemented for batch-processing many samples in a terminal environment. We introduce this system as *idCOV*.

## Pipeline

### Input

The pipeline (Figure 1) starts with the sequencing data derived from SARS-Cov-2 isolates in the format of pair-end FastQ files. These files are either available on the server or uploaded by the user through the web or the command-line interface. The fragments in the FastQ files are then aligned to a reference genome.

**Figure 1:**
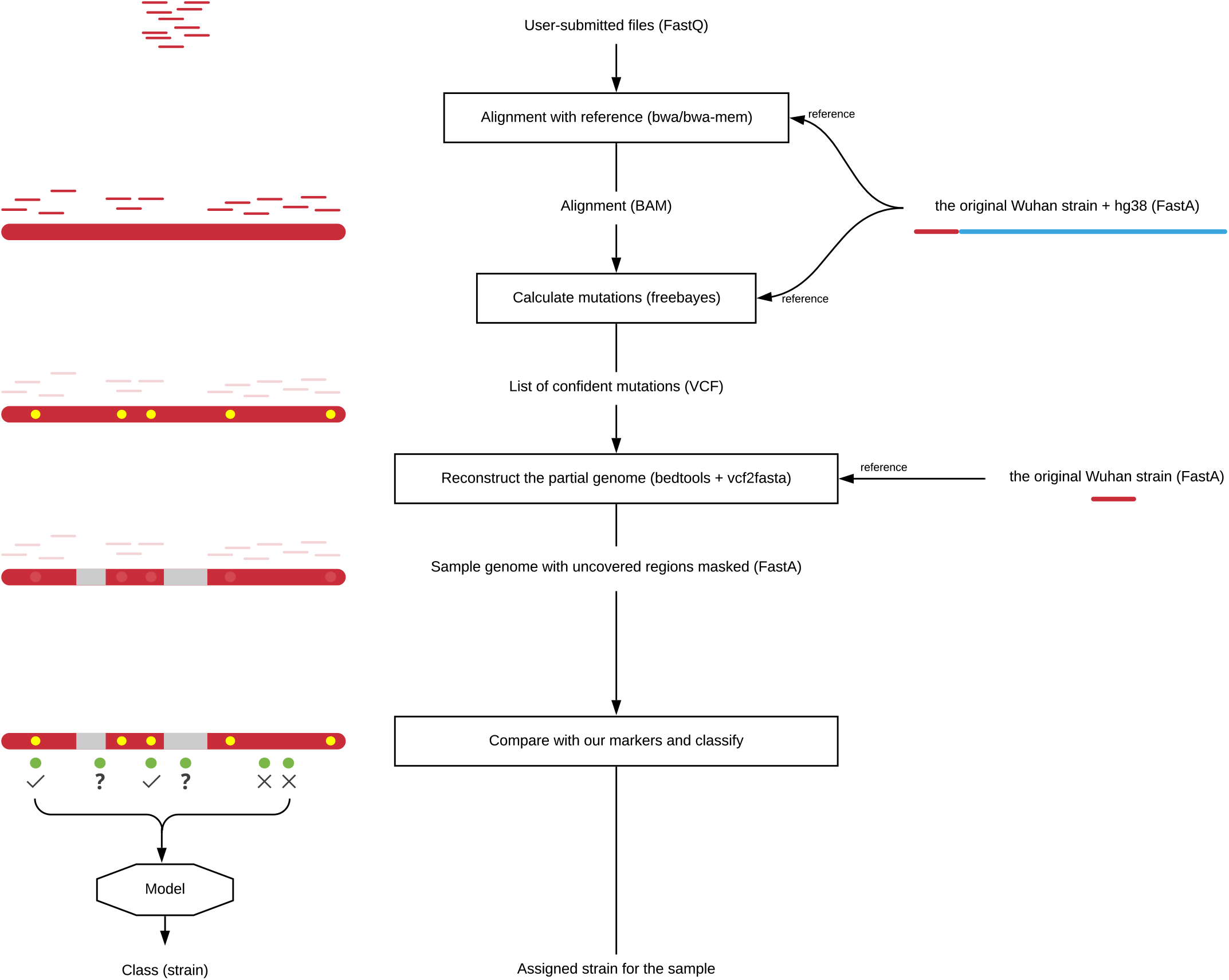
The bioinformatics pipeline of idCOV.

**Figure 2:**
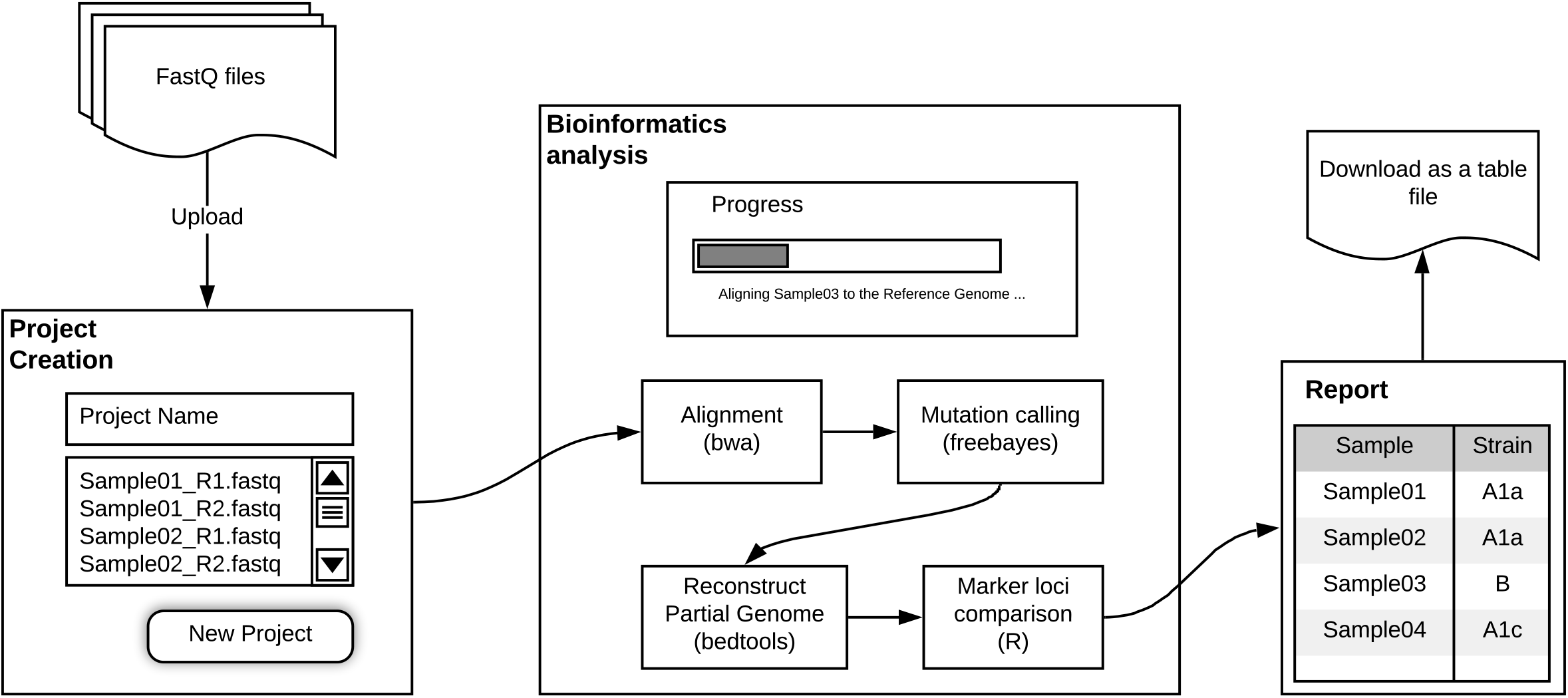
idCOV user experience.

### Alignment

Normally a virus reference genome is used for alignment. However, since sequencing library preparations might be generated from clinical material, some of the sequencing reads in the FastQ files may be from the human host. To account for this, we aggregated the Wuhan-Hu-1 genome (Wu, et al., 2020) with the GRCh38 human genome and used the combined genome as the reference for the alignment. The alignment is done using BWA, version 0.7.17 (Li and Durbin, 2009). The “bwa mem” algorithm is used by default. After the alignment, the result BAM files are first sorted by the fragment locations and then indexed, both using the SAMtools software, version 1.10 (Li, et al., 2009). This post-processing is to improve the performance of subsequent steps.

### Mutations Calculation

The virus reference genome (Wuhan-Hu-1) is seen as the original genome and any deviation from it is seen as a mutation. We run FreeBayes, version 1.3.2 (Garrison and Marth, 2012) on the BAM files generated above and the reference genome to calculate the mutations. The ploidy is set to one and the fragments mapping to the human genome are removed in the input to FreeBayes. The default values are used for the rest of the FreeBayes parameters. The output from FreeBayes is a list of confident mutations stored in a VCF file.

### Partial genome reconstruction

Depending on the depth and distribution of the sequencing, there might be regions in the viral genome to which no fragments were mapped. In the VCF output from FreeBayes, an absence of mutations due to unmapped regions must be distinguished from an actual absence of mutations (as deemed by FreeBayes.) Therefore, before we compare the markers, we reconstruct the partial genome of the isolate by masking the un-mapped regions in the reference gnome, and then apply the mutations as called by FreeBayes to the masked genome.

### Isolate identification

A marker profile is generated from the reconstructed genome. The marker profile consists of the base pairs at a list of predefined marker locations. These marker locations are dictated by the clade system. Currently, three clade systems are used in the classification step: the old Nextstrain, the new Nextstrain (Hodcroft, et al., 2020), and the GISAID clade system (e.V., 2020).

Finally, given a clade system, the marker profile is compared with the reference clade-defining marker profiles (Supplementary Table 1). If the marker profile of the isolate is the same as that of a designated clade, the isolate is called as the said clade. If the marker profile of the isolate is different from that of any of the clades, the clade with a marker profile most similar (in terms of Manhattan distance) is reported, along with the Manhattan distance between the two. When multiple clades are tied as having the least Manhattan distance, the clade of the isolate is considered ambiguous between the tied clades. If none of the Manhattan distances is below 4.5, the clade of the isolate is considered undetermined.

For two marker profiles *A* = (*a*_1_, *a*_2_, …, *a_n_*) and *B* = (*b*_1_, *b*_2_, …, *b*_n_), their Manhattan distance is defined as 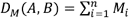, where *M_i_* = 0 when *a_i_* = *b_i_*, *M_i_* = 1 when *a_i_* ≠ *b_i_*, and *M_i_* = 0.5 when either *a_i_* or *b_i_* is undetermined due to lack of mapped reads at the marker region.

## Software

### The web user interface

After idCOV is deployed on a Linux environment, it can be accessed using any modern browser. A user can either upload the gzipped FastQ files stored on the client machine or specify a list of gzipped FastQ files already on the server file pool. Each pair of gzipped FastQ files needs to be named “*R1.fastq.gz” and “*R2.fastq.gz” so that their parity can be determined. A project can be created with the uploaded or specified files. The pipeline can be started by running a project. The sessions are tracked on the server, so the progress of any running project will not be lost when the client browser is closed. (Supplementary Figure 1)

### The command line interface

A command line interface will also be provided on the server-side after idCOV is deployed. Assuming terminal access (either direct or through SSH), a user can use the command “idcov” in the command line to run the pipeline directly on any gzipped FastQ files on the server.

### Data security

When deployed on a private computer or a cloud VM, all uploaded data is stored in a folder specified by a Docker volume binding parameter. The administrator of the computer can set the appropriate access permissions for this folder to ensure the data security.

### Implementation

The workflow management is done by using the Nextflow system (Di Tommaso, et al., 2017). The individual steps are broken down into Nextflow processes, and the deployment on multiple platforms is done by preparing a configuration for HPC and another for cloud environment. The latter can also be used on personal computers. A Datahike database (Kühne, et al., 2020) is used for storing user accounts, sessions, and job tracking. The server is written in Clojure using the Ring library. The web frontend is written in ClojureScript using the Fulcro library. The command-line interface is written in Python.

## Testing

The Clojure/ClojureScript code is unit-tested using the clojure.test library. The web UI is tested in Google Chrome Version 84.0.4147.105 (Official Build) (64-bit) and Mozilla Firefox Version 79.0.

We tested the entire workflow on 18 representative samples from a SARS-Cov-2 genomic sequencing dataset reported by the Peter Doherty Institute for Infection and Immunity and the Victorian Department of Health and Human Services in Australia (Seemann, et al., 2020). These samples included viral sequences representative of L, V, S, G, GR, and GH clades (GISAID clade nomenclature). idCOV correctly identified the clade of all viruses, regardless of the clade-defining marker profile used (i.e., Nextstrain old, new, and GISAID) (Figure 3). The full report is attached in the Supplementary Materials.

**Figure 3:**
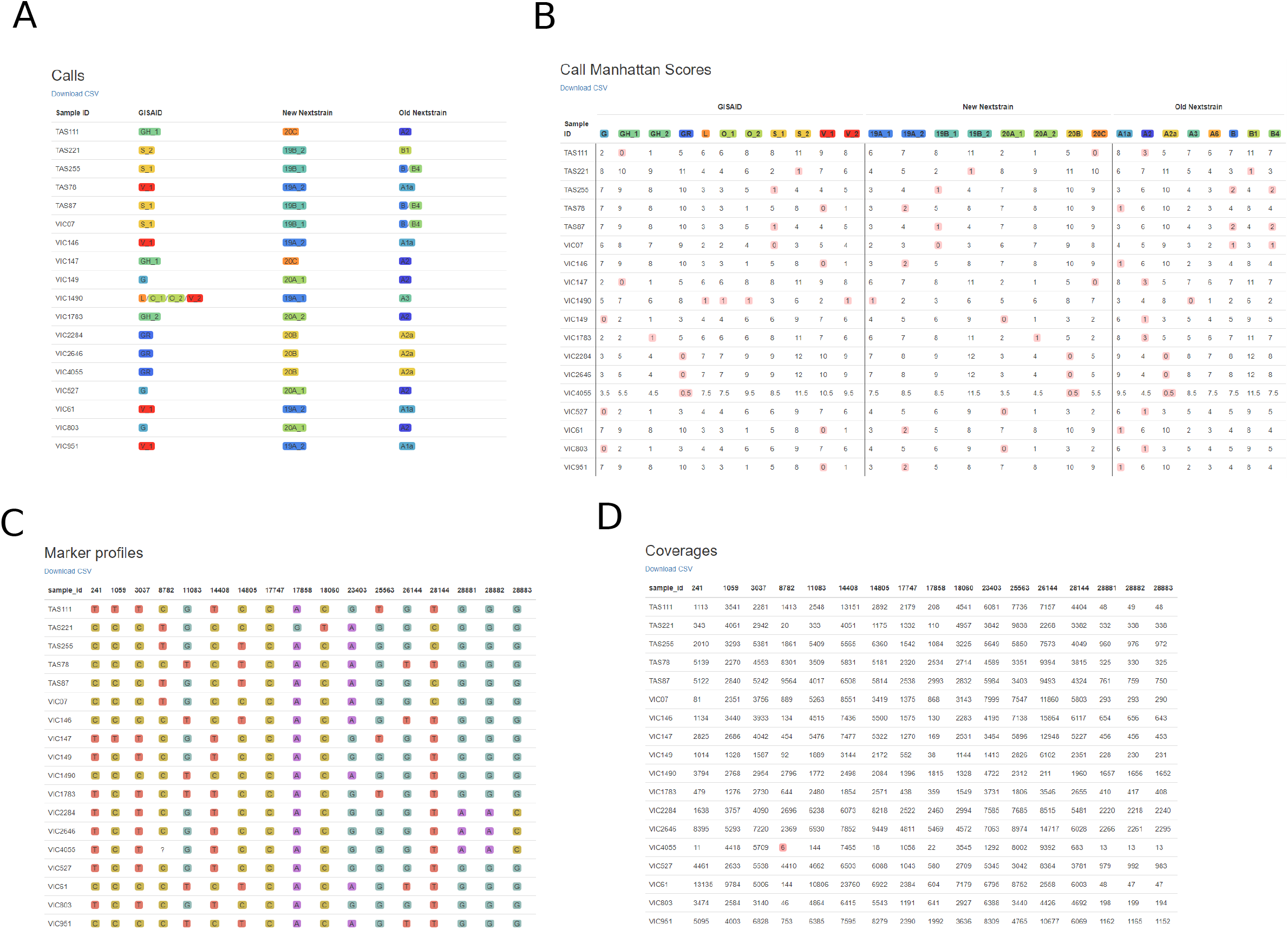
Screenshots of an example report of idCOV. The dataset consists of a random selection of isolates from Seemann, et al., 2020 covering all major clades. Table A) shows the final calls with color scheme consistent with Nextstrain.org. Table B) shows the Manhattan distances between every sample and every clade. A smaller Manhattan distance indicates a higher degree of similarity. Table C) shows the marker profiles and Table D) shows the marker coverages. When a marker is covered by fewer than 10 reads, the marker is deemed as undetermined and denoted as a question mark (?).

## Conclusion

Rapid classification of SARS-Cov-2 viruses into their respective genetic clades helps characterize transmission patterns and assess local and global epidemiologic trends. idCOV provides an easy portal to a complete bioinformatics pipeline that identifies a virus’ clade using the three most widely used clade systems. The system is internally tested and has been shown to provide valuable insights to clinicians and biologists in a timely manner.

## References

CSSE. COVID-19 Dashboard by the Center for Systems Science and Engineering (CSSE) at Johns Hopkins University (JHU). In. ArcGIS Online (arcgis.com); 2020.

Di Tommaso, P., et al. Nextflow enables reproducible computational workflows. Nature biotechnology 2017;35(4):316–319.

Dong, E., Du, H. and Gardner, L. An interactive web-based dashboard to track COVID-19 in real time. The Lancet infectious diseases 2020;20(5):533–534.

e.V., F.v.G. GISAID - Clade and lineage nomenclature aids in genomic epidemiology of active hCoV-19 viruses. In. Gisaid.org: GISAID; 2020.

Garrison, E. and Marth, G. Haplotype-based variant detection from short-read sequencing. arXiv preprint arXiv:1207.3907 2012.

Gonzalez-Reiche, A.S., et al. Introductions and early spread of SARS-CoV-2 in the New York City area. Science 2020.

Hodcroft, E.B., et al. Year-letter Genetic Clade Naming for SARS-CoV-2 on Nextstain.org. In. Nextstrain.org: Nextstrain; 2020.

Kühne, K., Weilbach, C. and Prokopov, N. Datahike is a durable Datalog database powered by an efficient Datalog query engine. In. Github.com; 2020.

Lescure, F.-X., et al. Clinical and virological data of the first cases of COVID-19 in Europe: a case series. The Lancet Infectious Diseases 2020.

Li, H. and Durbin, R. Fast and accurate short read alignment with Burrows–Wheeler transform. bioinformatics 2009;25(14):1754–1760.

Li, H., et al. The sequence alignment/map format and SAMtools. Bioinformatics 2009;25(16):2078–2079.

Seemann, T., et al. Tracking the COVID-19 pandemic in Australia using genomics. medRxiv 2020.

Wu, F., et al. A new coronavirus associated with human respiratory disease in China. Nature 2020;579(7798):265–269.

