## Supplementary Material 1 for "idCOV: a pipeline for quick clade identification of SARS-CoV-2 isolates": index.html

idCOV Report


### idCOV Report

#### Calls

Download CSV

| Sample ID | GISAID | New Nextstrain | Old Nextstrain |
| --- | --- | --- | --- |
| TAS111 | GH\_1 | 20C | A2 |
| TAS221 | S\_2 | 19B\_2 | B1 |
| TAS255 | S\_1 | 19B\_1 | B/B4 |
| TAS78 | V\_1 | 19A\_2 | A1a |
| TAS87 | S\_1 | 19B\_1 | B/B4 |
| VIC07 | S\_1 | 19B\_1 | B/B4 |
| VIC146 | V\_1 | 19A\_2 | A1a |
| VIC147 | GH\_1 | 20C | A2 |
| VIC149 | G | 20A\_1 | A2 |
| VIC1490 | L/O\_1/O\_2/V\_2 | 19A\_1 | A3 |
| VIC1783 | GH\_2 | 20A\_2 | A2 |
| VIC2284 | GR | 20B | A2a |
| VIC2646 | GR | 20B | A2a |
| VIC4055 | GR | 20B | A2a |
| VIC527 | G | 20A\_1 | A2 |
| VIC61 | V\_1 | 19A\_2 | A1a |
| VIC803 | G | 20A\_1 | A2 |
| VIC951 | V\_1 | 19A\_2 | A1a |

#### Call Manhattan Scores

The number in each cell is the Manhattan distance between the sample and the clade. The lower the Manhattan distance, the more similar between the marker profiles. The clade(s) with the lowest manhattan distance is highlighted. The highlighted clade(s) are regarded as the assigned clade for the sample.

Download CSV

|  | GISAID | | | | | | | | | | | New Nextstrain | | | | | | | | Old Nextstrain | | | | | | | |
| --- | --- | --- | --- | --- | --- | --- | --- | --- | --- | --- | --- | --- | --- | --- | --- | --- | --- | --- | --- | --- | --- | --- | --- | --- | --- | --- | --- |
| Sample ID | G | GH\_1 | GH\_2 | GR | L | O\_1 | O\_2 | S\_1 | S\_2 | V\_1 | V\_2 | 19A\_1 | 19A\_2 | 19B\_1 | 19B\_2 | 20A\_1 | 20A\_2 | 20B | 20C | A1a | A2 | A2a | A3 | A6 | B | B1 | B4 |
| TAS111 | 2 | 0 | 1 | 5 | 6 | 6 | 8 | 8 | 11 | 9 | 8 | 6 | 7 | 8 | 11 | 2 | 1 | 5 | 0 | 8 | 3 | 5 | 7 | 6 | 7 | 11 | 7 |
| TAS221 | 8 | 10 | 9 | 11 | 4 | 4 | 6 | 2 | 1 | 7 | 6 | 4 | 5 | 2 | 1 | 8 | 9 | 11 | 10 | 6 | 7 | 11 | 5 | 4 | 3 | 1 | 3 |
| TAS255 | 7 | 9 | 8 | 10 | 3 | 3 | 5 | 1 | 4 | 4 | 5 | 3 | 4 | 1 | 4 | 7 | 8 | 10 | 9 | 3 | 6 | 10 | 4 | 3 | 2 | 4 | 2 |
| TAS78 | 7 | 9 | 8 | 10 | 3 | 3 | 1 | 5 | 8 | 0 | 1 | 3 | 2 | 5 | 8 | 7 | 8 | 10 | 9 | 1 | 6 | 10 | 2 | 3 | 4 | 8 | 4 |
| TAS87 | 7 | 9 | 8 | 10 | 3 | 3 | 5 | 1 | 4 | 4 | 5 | 3 | 4 | 1 | 4 | 7 | 8 | 10 | 9 | 3 | 6 | 10 | 4 | 3 | 2 | 4 | 2 |
| VIC07 | 6 | 8 | 7 | 9 | 2 | 2 | 4 | 0 | 3 | 5 | 4 | 2 | 3 | 0 | 3 | 6 | 7 | 9 | 8 | 4 | 5 | 9 | 3 | 2 | 1 | 3 | 1 |
| VIC146 | 7 | 9 | 8 | 10 | 3 | 3 | 1 | 5 | 8 | 0 | 1 | 3 | 2 | 5 | 8 | 7 | 8 | 10 | 9 | 1 | 6 | 10 | 2 | 3 | 4 | 8 | 4 |
| VIC147 | 2 | 0 | 1 | 5 | 6 | 6 | 8 | 8 | 11 | 9 | 8 | 6 | 7 | 8 | 11 | 2 | 1 | 5 | 0 | 8 | 3 | 5 | 7 | 6 | 7 | 11 | 7 |
| VIC1490 | 5 | 7 | 6 | 8 | 1 | 1 | 1 | 3 | 6 | 2 | 1 | 1 | 2 | 3 | 6 | 5 | 6 | 8 | 7 | 3 | 4 | 8 | 0 | 1 | 2 | 6 | 2 |
| VIC149 | 0 | 2 | 1 | 3 | 4 | 4 | 6 | 6 | 9 | 7 | 6 | 4 | 5 | 6 | 9 | 0 | 1 | 3 | 2 | 6 | 1 | 3 | 5 | 4 | 5 | 9 | 5 |
| VIC1783 | 2 | 2 | 1 | 5 | 6 | 6 | 6 | 8 | 11 | 7 | 6 | 6 | 7 | 8 | 11 | 2 | 1 | 5 | 2 | 8 | 3 | 5 | 5 | 6 | 7 | 11 | 7 |
| VIC2284 | 3 | 5 | 4 | 0 | 7 | 7 | 9 | 9 | 12 | 10 | 9 | 7 | 8 | 9 | 12 | 3 | 4 | 0 | 5 | 9 | 4 | 0 | 8 | 7 | 8 | 12 | 8 |
| VIC2646 | 3 | 5 | 4 | 0 | 7 | 7 | 9 | 9 | 12 | 10 | 9 | 7 | 8 | 9 | 12 | 3 | 4 | 0 | 5 | 9 | 4 | 0 | 8 | 7 | 8 | 12 | 8 |
| VIC4055 | 3.5 | 5.5 | 4.5 | 0.5 | 7.5 | 7.5 | 9.5 | 8.5 | 11.5 | 10.5 | 9.5 | 7.5 | 8.5 | 8.5 | 11.5 | 3.5 | 4.5 | 0.5 | 5.5 | 9.5 | 4.5 | 0.5 | 8.5 | 7.5 | 7.5 | 11.5 | 7.5 |
| VIC527 | 0 | 2 | 1 | 3 | 4 | 4 | 6 | 6 | 9 | 7 | 6 | 4 | 5 | 6 | 9 | 0 | 1 | 3 | 2 | 6 | 1 | 3 | 5 | 4 | 5 | 9 | 5 |
| VIC61 | 7 | 9 | 8 | 10 | 3 | 3 | 1 | 5 | 8 | 0 | 1 | 3 | 2 | 5 | 8 | 7 | 8 | 10 | 9 | 1 | 6 | 10 | 2 | 3 | 4 | 8 | 4 |
| VIC803 | 0 | 2 | 1 | 3 | 4 | 4 | 6 | 6 | 9 | 7 | 6 | 4 | 5 | 6 | 9 | 0 | 1 | 3 | 2 | 6 | 1 | 3 | 5 | 4 | 5 | 9 | 5 |
| VIC951 | 7 | 9 | 8 | 10 | 3 | 3 | 1 | 5 | 8 | 0 | 1 | 3 | 2 | 5 | 8 | 7 | 8 | 10 | 9 | 1 | 6 | 10 | 2 | 3 | 4 | 8 | 4 |

#### Phylogeny

##### GISAID

##### Old Nextstrain

##### New Nextstrain

#### Marker profiles

Download CSV

| sample\_id | 241 | 1059 | 3037 | 8782 | 11083 | 14408 | 14805 | 17747 | 17858 | 18060 | 23403 | 25563 | 26144 | 28144 | 28881 | 28882 | 28883 |
| --- | --- | --- | --- | --- | --- | --- | --- | --- | --- | --- | --- | --- | --- | --- | --- | --- | --- |
| TAS111 | T | T | T | C | G | T | C | C | A | C | G | T | G | T | G | G | G |
| TAS221 | C | C | C | T | G | C | C | C | G | T | A | G | G | C | G | G | G |
| TAS255 | C | C | C | T | G | C | T | C | A | C | A | G | G | C | G | G | G |
| TAS78 | C | C | C | C | T | C | T | C | A | C | A | G | T | T | G | G | G |
| TAS87 | C | C | C | T | G | C | T | C | A | C | A | G | G | C | G | G | G |
| VIC07 | C | C | C | T | G | C | C | C | A | C | A | G | G | C | G | G | G |
| VIC146 | C | C | C | C | T | C | T | C | A | C | A | G | T | T | G | G | G |
| VIC147 | T | T | T | C | G | T | C | C | A | C | G | T | G | T | G | G | G |
| VIC149 | T | C | T | C | G | T | C | C | A | C | G | G | G | T | G | G | G |
| VIC1490 | C | C | C | C | T | C | C | C | A | C | A | G | G | T | G | G | G |
| VIC1783 | T | C | T | C | T | T | C | C | A | C | G | T | G | T | G | G | G |
| VIC2284 | T | C | T | C | G | T | C | C | A | C | G | G | G | T | A | A | C |
| VIC2646 | T | C | T | C | G | T | C | C | A | C | G | G | G | T | A | A | C |
| VIC4055 | T | C | T | Z | G | T | C | C | A | C | G | G | G | T | A | A | C |
| VIC527 | T | C | T | C | G | T | C | C | A | C | G | G | G | T | G | G | G |
| VIC61 | C | C | C | C | T | C | T | C | A | C | A | G | T | T | G | G | G |
| VIC803 | T | C | T | C | G | T | C | C | A | C | G | G | G | T | G | G | G |
| VIC951 | C | C | C | C | T | C | T | C | A | C | A | G | T | T | G | G | G |

#### Coverages

Download CSV

| sample\_id | 241 | 1059 | 3037 | 8782 | 11083 | 14408 | 14805 | 17747 | 17858 | 18060 | 23403 | 25563 | 26144 | 28144 | 28881 | 28882 | 28883 |
| --- | --- | --- | --- | --- | --- | --- | --- | --- | --- | --- | --- | --- | --- | --- | --- | --- | --- |
| TAS111 | 1113 | 3541 | 2281 | 1413 | 2548 | 13151 | 2892 | 2179 | 208 | 4541 | 6081 | 7736 | 7157 | 4404 | 48 | 49 | 48 |
| TAS221 | 343 | 4061 | 2942 | 20 | 333 | 4051 | 1175 | 1332 | 110 | 4957 | 3842 | 9838 | 2268 | 3382 | 332 | 338 | 338 |
| TAS255 | 2010 | 3293 | 5381 | 1861 | 5409 | 5565 | 6360 | 1542 | 1084 | 3225 | 5649 | 5850 | 7573 | 4049 | 960 | 976 | 972 |
| TAS78 | 5139 | 2270 | 4553 | 8301 | 3509 | 5831 | 5181 | 2320 | 2534 | 2714 | 4589 | 3351 | 9394 | 3815 | 325 | 330 | 325 |
| TAS87 | 5122 | 2840 | 5242 | 9564 | 4017 | 6508 | 5814 | 2538 | 2993 | 2832 | 5984 | 3403 | 9493 | 4324 | 761 | 759 | 750 |
| VIC07 | 81 | 2351 | 3756 | 889 | 5263 | 8551 | 3419 | 1375 | 868 | 3143 | 7999 | 7547 | 11860 | 5803 | 293 | 293 | 290 |
| VIC146 | 1134 | 3440 | 3933 | 134 | 4515 | 7436 | 5500 | 1575 | 130 | 2283 | 4195 | 7138 | 15864 | 6117 | 654 | 656 | 643 |
| VIC147 | 2825 | 2686 | 4042 | 454 | 5476 | 7477 | 5322 | 1270 | 169 | 2531 | 3464 | 5896 | 12948 | 5227 | 456 | 456 | 453 |
| VIC149 | 1014 | 1328 | 1587 | 92 | 1889 | 3144 | 2172 | 552 | 38 | 1144 | 1413 | 2826 | 6102 | 2351 | 228 | 230 | 231 |
| VIC1490 | 3794 | 2768 | 2954 | 2796 | 1772 | 2498 | 2084 | 1396 | 1815 | 1328 | 4722 | 2312 | 211 | 1960 | 1657 | 1656 | 1652 |
| VIC1783 | 479 | 1276 | 2730 | 644 | 2480 | 1854 | 2571 | 438 | 359 | 1549 | 3731 | 1806 | 3546 | 2655 | 410 | 417 | 408 |
| VIC2284 | 1638 | 3757 | 4090 | 2696 | 6238 | 6073 | 8218 | 2522 | 2460 | 2994 | 7585 | 7685 | 8515 | 5481 | 2220 | 2218 | 2240 |
| VIC2646 | 8395 | 5293 | 7220 | 2369 | 6930 | 7852 | 9449 | 4811 | 5469 | 4572 | 7063 | 8974 | 14717 | 6028 | 2266 | 2261 | 2295 |
| VIC4055 | 11 | 4418 | 5709 | 6 | 144 | 7465 | 18 | 1058 | 22 | 3545 | 1292 | 8002 | 9392 | 683 | 13 | 13 | 13 |
| VIC527 | 4461 | 2633 | 5538 | 4410 | 4662 | 6503 | 6088 | 1043 | 580 | 2709 | 5345 | 3042 | 8384 | 3781 | 979 | 992 | 983 |
| VIC61 | 13135 | 9784 | 5006 | 144 | 10806 | 23760 | 6922 | 2384 | 604 | 7179 | 6795 | 8752 | 2558 | 6003 | 48 | 47 | 47 |
| VIC803 | 3474 | 2584 | 3140 | 46 | 4864 | 6415 | 5543 | 1191 | 641 | 2927 | 6388 | 3440 | 4426 | 4692 | 198 | 199 | 194 |
| VIC951 | 5095 | 4003 | 6828 | 753 | 6385 | 7595 | 8279 | 2390 | 1992 | 3636 | 8309 | 4765 | 10677 | 6069 | 1162 | 1165 | 1152 |
